# Genome analysis of Parmales, a sister group of diatoms, reveals the evolutionary specialization of diatoms from phago-mixotrophs to photoautotrophs

**DOI:** 10.1101/2022.09.09.507052

**Authors:** Hiroki Ban, Shinya Sato, Shinya Yoshikawa, Kazumasa Yamada, Yoji Nakamura, Mutsuo Ichinomiya, Naoki Sato, Romain Blanc-Mathieu, Hisashi Endo, Akira Kuwata, Hiroyuki Ogata

## Abstract

The order Parmales (Bolidophyceae) is a minor group of pico-sized eukaryotic marine phytoplankton that contains species with cells surrounded by silica plates. Previous studies revealed that Parmales is a member of ochrophytes and sister to diatoms (Bacillariophyta), the most successful phytoplankton group in the modern ocean. Therefore, parmalean genomes can serve as a reference to elucidate both the evolutionary events that differentiated these two lineages and the genomic basis for the ecological success of diatoms vs. the more cryptic lifestyle of parmaleans. Here, we compared the genomes of eight parmaleans and five diatoms to explore their physiological and evolutionary differences. Parmaleans were predicted to be phago-mixotrophs. By contrast, diatoms have undergone loss of genes related to phagocytosis, indicating the ecological specialization from phago-mixotroph to photoautotroph in the early evolution of diatoms. Furthermore, diatoms showed significant enrichment in gene sets involved in silica metabolism, nutrient uptake capacity, carbon concentrating mechanisms, and iron uptake in comparison with parmaleans. Overall, our results suggest a strong evolutionary link between the loss of phago-mixotrophy and specialization to a silicified photoautotrophic life stage early in diatom evolution after diverging from the Parmales lineage.

## Introduction

Parmales is a group of pico-sized (2–5 μm) eukaryotic marine phytoplankton with cells surrounded by silicified plates^1^. Parmaleans are widespread in the ocean, from polar to subtropical regions, and are relatively abundant in polar and subarctic regions. Parmalean sequences are most abundant in the picoplanktonic fraction (0.8–5 μm) of the global ocean metabarcoding data from *Tara* Oceans and represent at most 4% of the sequences of photosynthetic organisms and less than 1% on average^2^. Currently, only 17 taxa of parmaleans have been described^3,4^. SEM and TEM observations, molecular phylogenetics, and photosynthetic pigment analyses indicated that Parmales belongs to ochrophytes (class Bolidophyceae)^5^ and is the sister taxon of diatoms (phylum Bacillariophyta). Bolidophyceae also contains pico-sized photosynthetic naked flagellates (called bolidomonads) that mainly inhabit subtropical waters^6^. Recent phylogenetic analyses using several newly isolated strains revealed that flagellated bolidomonad species belong to the silicified and non-flagellated parmalean genus *Triparma* within Bolidophyceae, suggesting that the *Triparma* life cycle switches between silicified/non-flagellated and naked/flagellated stages^2^.

Diatoms are the most successful phytoplankton group in the modern ocean; they have high diversity (ca. 10^5^ species^7^) and high primary productivity, contributing an estimated 20% of photosynthesis on Earth. Diatoms are thought to be particularly successful in dynamic environments such as upwelling areas, and it has been suggested that their ecological success is supported by traits such as silicified cell wall defense^8^ and luxury nutrient uptake^9^. However, despite advances in understanding diatom genomes during the last two decades, the reasons underlying the success of diatoms in modern oceans remain poorly understood. To understand the ecological success of diatoms, characterization of the evolution of physiology-related genes in this taxon is necessary.

Although parmaleans are the closest relatives of diatoms, they show much lower biomass, species diversity, and ecological impact than their sister taxon. The proposed parmalean life cycle, which switches between silicified/non-flagellated and naked/flagellated stages, is similar to the proposed origin of diatoms^2^. Ancestral diatoms were possibly haploid flagellates that formed silicified diploid zygotes^10^. The mitotic division of the zygote might have taken place preferentially to give rise to centric diatoms^11^, which is the most ancient diatom group with a diploid vegetative stage producing naked flagellated haploid male gametes for sexual reproduction^12^. Thus, a comparison of parmaleans and diatoms is expected to provide important clues on differences in their ecological strategies and evolutionary paths. To date, only limited genomic data on parmaleans have been available^13^, and the genomic features and evolutionary events that led to differences between them and diatoms have remained unstudied. In this study, we generated seven novel parmalean genome assemblies. These seven draft genomes, one previously determined parmalean genome, and five publicly available diatom genomes were used to perform a comparative genome analysis. Our results delineate the evolutionary trajectories of these two lineages after their divergence and correlate their differential ecological features with their genomic functions.

## Results and Discussion

### General genomic features

In this study, we obtained whole-genome sequences of seven parmaleans, including six strains from two genera (*Triparma* and *Tetraparma*) that are frequently observed in the subarctic Pacific Ocean^4,14^, as well as one strain from an undescribed taxon that is phylogenetically and morphologically distinct from known parmaleans (hereafter referred to as ‘Scaly parma’, Sato et al. in prep.). Together with the previously sequenced *Triparma laevis* f. *inornata* genome^13^, we built a database of eight parmalean strain genomes. The parmalean genomes were similar in size, ranging from 31.0 Mb for ‘Scaly parma’ to 43.6 Mb for *Tetraparma gracilis* (Table 1). The predicted numbers of genes ranged from 12,177 for ‘Scaly parma’ to 16,002 for *Triparma laevis* f. *longispina* (Table 1). These genome sizes are relatively constant compared to diatom genomes and similar to those of *Thalassiosira pseudonana* (32.4 Mb)^15^ and *Phaeodactylum tricornutum* (27.4 Mb)^16^, which have rather small genomes among diatoms.

**Table 1.** Assembly and annotation results and statistics

|  | *Triparma*<br>*laevis* f. *inornata* | *Triparma*<br>*laevis* f. *longispina* | *Triparma*<br>*verrucosa* | *Triparma*<br>*strigata* | *Triparma*<br>*retinervis* | *Triparma*<br>*colmacea* | *Tetraparma*<br>*gracilis* | Scaly<br>parma |
| --- | --- | --- | --- | --- | --- | --- | --- | --- |
| Genome<br>size (Mbp) | 42.6 | 41.4 | 35.5 | 35.2 | 36.5 | 43.0 | 43.6 | 31.0 |
| No. of<br>scaffolds | 902 | 1,055 | 659 | 634 | 8,760 | 1,858 | 7,082 | 1,921 |
| N50<br>(kbp) | 83.2 | 77.9 | 74.0 | 73.4 | 8.2 | 63.5 | 10.7 | 51.7 |
| GC (%) | 51.3 | 51.1 | 52.1 | 52.2 | 52.4 | 51.0 | 64.8 | 51.0 |
| No. of<br>predicted protein-coding genes | 13,396 | 16,002 | 14,488 | 14,364 | 13,636 | 13,919 | 15,310 | 12,177 |
| BUSCO<br>Complete genes (%) | 74 | 93 | 94 | 94 | 70 | 92 | 65 | 93 |

Analysis of orthologous groups (OGs) revealed 62,363 OGs among the parmaleans (8 strains), diatoms (5 strains), and other stramenopiles (5 strains). Phylogenomic analysis based on 164 single-copy OGs clearly shows parmaleans are monophyletic and sister to diatoms (Fig. 1a). 34,292 OGs were present only in diatoms or parmaleans and not in other stramenopiles (Fig. 1b: yellow + orange + purple + green in diatoms and Parmales). Of those, only 1,448 OGs were shared by diatoms and parmaleans (Fig. 1b: yellow). 20,974 OGs were specific to diatoms (diatom-specific OGs, Fig. 1b: orange + green in diatoms), and 11,870 OGs were specific to parmaleans (Parmales-specific OGs, Fig. 1b: purple and green in Parmales). 99.7% of the genes in the core OGs conserved in all analysed strains (1,153 OGs, Fig. 1b: red) had InterPro domains, and 77.1% of the genes in the OGs shared only by diatoms and parmaleans (1,448 OGs, Fig. 1b: yellow) had InterPro domains. By contrast, only 24.9% of genes in diatom-specific OGs (20,974 OGs, Fig. 1b: orange + green in diatoms) and 55.1% of genes in parmalean-specific OGs (11,870 OGs, Fig. 1b: purple and green in Parmales) had InterPro domains. These results reveal that many lineage-specific genes are functionally unknown.

**Fig. 1.**
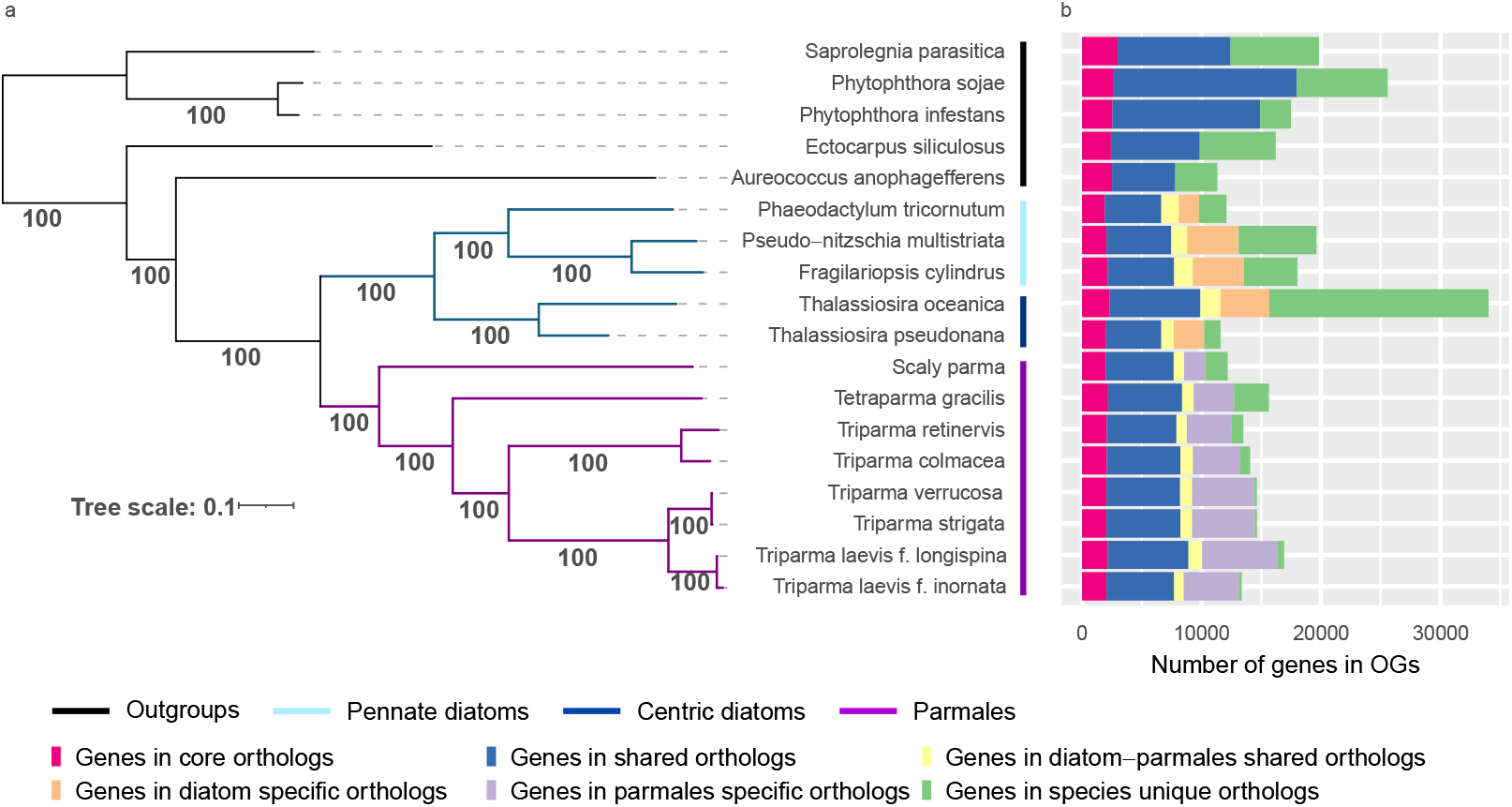
Phylogenetic relationships of diatoms, parmaleans (Parmales), and stramenopiles, and number of shared genes in OGs. (a) Maximum likelihood tree estimated by RAxML with 164 single-copy OGs. Blue and purple branches are diatom and Parmales clades, respectively. The numbers on the branches represent bootstrap values. The coloured bars indicate the group where each taxon belongs (black: outgroup, light blue: pennate diatoms, deep blue: centric diatoms, purple: Parmales). (b) The barplot represents the groups genes inferred through orthologous gene clustering.

### Differentially enriched protein domains

By comparing the eight parmalean and five diatom genomes, we found 64 and 315 InterPro domains in which the diatom and Parmales lineages, respectively, were significantly enriched. Cyclin domains and heat-shock transcription factor domains in which diatoms are enriched have been known to exhibit expanded gene families compared with other eukaryotes^16,17^ (Fig. 2a). In addition, diatoms were enriched in protease domains and sulfotransferase domains. Serine proteases and metalloproteases are known to be induced by limitations of nitrogen, iron, silicon, and light^18,19^. Sulfotransferase is an enzyme that catalyses sulfonation and is thought to be related to programmed cell death in *Skeletonema marinoi*, a bloom-forming marine diatom^20^. These gene families are thought to be related to the stress response process in diatoms.

**Fig. 2.**
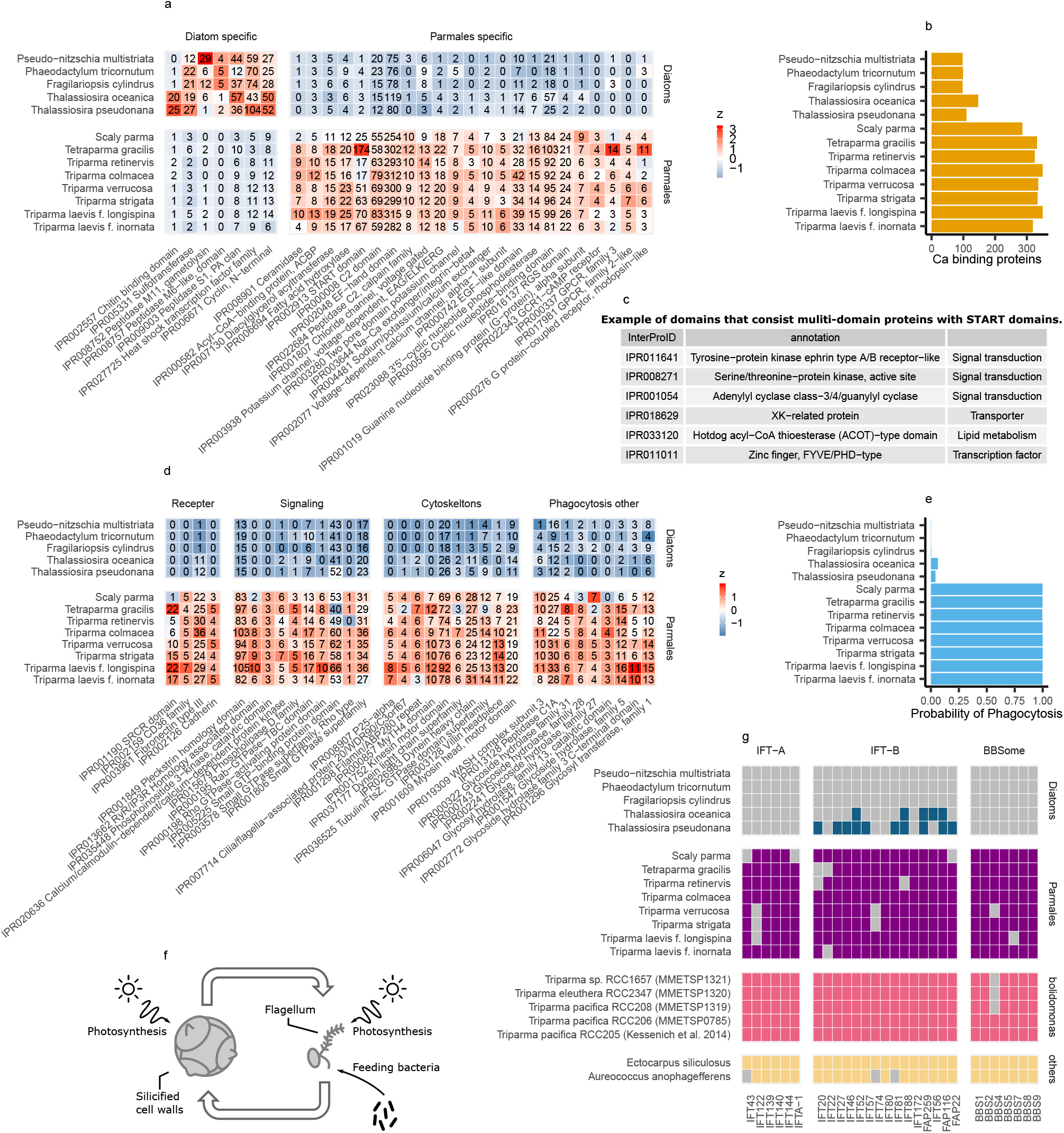
(a) Diatom and Parmales genomes enriched in InterPro domains. We manually clustered and selected the domains that appear to be involved in a specific process. The colours are scaled in ascending order from blue to red by the z-value in each row. (b) Number of genes annotated with GO:0005516 (calcium ion binding) by InterProScan. (c) An example of InterPro domains composed of multi-domain proteins including START domains. (d) Parmalean genomes enriched in InterPro domains thought to be related to phagocytosis. The colours are scaled in ascending order from blue to red by the z-value in each row. InterPro domains that were not statistically significant but were considered important are marked with an asterisk (*). (e) Probability of phagotrophy predicted by the Burns et al. (2019) tool. (f) Schematic view of hypothesized parmalean life cycle. (g) Presence (filled square) or absence (or loss: grey square) of genes/transcripts related to intraflagellar transport (IFT) subunits.

InterPro domains in which parmaleans were enriched included those involved in intracellular signalling pathways, such as the G protein signalling, cyclic nucleotide signalling, calcium signalling, and action potential pathways (Fig. 2a). G protein-coupled receptors were involved in responses to sexual cues in the planktonic diatom *Pseudo-nitzschia multistriata*^17^, and to colonization in *Phaeodactylum tricornutum*, a planktonic diatom that also has a benthic morphotype^21^. Diatoms also exhibit action potential signalling to modulate their cellular motility^22,23^. Furthermore, parmaleans encode a strikingly greater number of calcium-binding proteins (up to 300) that could act as messenger molecules^24^ (Fig. 2b). Enrichment of intracellular signalling pathways in parmaleans may be associated with their putative alternating life cycle stages (i.e., silicified/non-flagellated and naked/flagellated cell stages^2^).

Parmalean genomes were notably enriched in domains associated with lipids and fatty acids (Fig. 2a). For example, diacylglycerol acyltransferase is an enzyme for the terminal step in the production of triacylglycerol, the main component of stored lipids^25^. The steroidogenic acute regulatory protein-related lipid transfer (START) domain that binds to lipids and sterols^26^ is one of the domains in which parmalean genomes are most enriched, with up to 174 domains in *Tetraparma gracilis*. This domain sometimes consists of multi-domain proteins and works in lipid trafficking, lipid metabolism, and cell signalling in animals and land plants^26^. START domain-containing proteins in parmaleans also contain other functional domains, such as lipid metabolism enzymes, transporters, kinases, and transcription factors (Fig. 2c). These results suggest diverse lipid-related physiological processes in parmaleans.

### Phagotrophy

Some InterPro domains in which parmaleans are enriched are known to be involved in phagotrophy^27^, including cell adhesion^28^, intercellular signalling (e.g., small GTP-binding proteins such as Rho)^29^, cytoskeleton^30^, lysosome^31^, and WASH^32^ (WASP and SCAR homolog) complex proteins (Fig. 2d). Specific genetic markers of phagotrophy are not known, but parmaleans were predicted as phago-mixotrophs (high scores > 0.98) according to a gene-based phago-mixotrophy prediction model^27^, whereas diatoms were not (low scores < 0.07) (Fig. 2e). This result suggests that parmaleans are capable of phagocytosis. We also applied this prediction model to the bolidomonads (naked/flagellated parmaleans) transcriptomes, and bolidomonads were also predicted as phago-mixotrophs (high scores > 0.98, Fig. 2e). Although there is no experimental evidence of phagocytosis in silicified parmaleans, a field study demonstrated that bolidomonads feed on cyanobacteria^33,34^. As transcriptome data reflect gene repertoires expressed under specific physiological conditions, bolidomonads might be phagotrophic. It remains unclear which life cycle stages of the parmaleans that we analysed are phagotrophic. However, assuming that bolidomonads indeed represent a part of the parmalean life cycle^3^, and a possibility that the silicified parmalean cell wall could physically interfere with feeding bacteria, it is likely that parmaleans perform phagocytosis in their putative naked/flagellated stage (Fig. 2f).

In the following sections, we move from the analysis of enriched domains to more focused investigation of genes in specific pathways and functions.

### Flagellum

To investigate the possibility that parmaleans can produce a flagellated cell^2^, we searched for genes responsible for flagellar motility (i.e., intraflagellar transport (IFT) subunit genes^35^ [IFT-A complex (6 genes), IFT-B complex (15 genes), BBSome (7 genes)]) in the parmalean and diatom genomes and bolidomonad transcriptomes. Flagellum structural genes for tubulin, radial spokes, dynein arms, and the central pair complex were excluded from analysis because these genes are also involved in other processes/structures (such as the centriole in *Triparma laevis*^36^) and are not unique to the flagellum. For this analysis, bolidomonad transcriptomes and centric diatom genomes were considered as positive controls because of the presence of the flagellar structure^6^ and the presence of flagellated sperm in their life cycle^37^, respectively. Similarly, pennate diatom genomes were considered as negative controls because flagellar structures have never been observed in this group, despite the accumulated knowledge concerning sexual reproduction^38^.

A nearly-full set of the flagellar genes were found in parmalean genomes and bolidomonad transcriptomes, whereas IFT-A and BBsome were completely absent in both types of diatoms (Fig. 2g). IFT-B was partially lost in centric diatoms and completely lost in pennate diatoms. These results suggest that parmaleans have a flagellated stage in their life cycle and are consistent with the idea that parmaleans are phago-mixotrophic in their putative naked/flagellated stage. Jensen et al.^39^ speculated that the two central microtubules were dispensed with in sperms of centric diatoms. Given the detection of the nearly-full set of flagellar genes in the parmaleans vs. the complete lack of IFT-A and BBSome and partial loss of IFT-B in the centric diatoms, it is possible that evolutionary pressure to maintain the flagellated stage is higher in parmaleans than in centric diatoms (e.g., because of the presence of a frequent or prolonged flagellated stage in parmaleans, which is not expected for sperms of centric diatoms).

### Nitrogen metabolism

The number of transporter genes involved in the uptake of nitrogen sources differed greatly between diatoms and parmaleans (Fig. 3a). Parmaleans had 1–3 nitrate/nitrite transporter genes, whereas diatoms had 4–9. Only one urea transporter gene was present in each parmalean, whereas 3–6 genes were present in each diatom. Diatoms tended to have more ammonia transporter genes than parmaleans, although the difference was not as obvious as for the other transporters (1–4 genes for parmaleans vs. 3–8 for diatoms). Genes for the vacuolar nitrate transporter, which stores nitrogen sources in the vacuole^40^, were absent from parmalean genomes. This suggests that parmaleans may be less competent to store nitrogen sources than diatoms, although the possibility that parmaleans encode another non-homologous vacuolar nitrate transporter is not excluded.

**Fig. 3.**
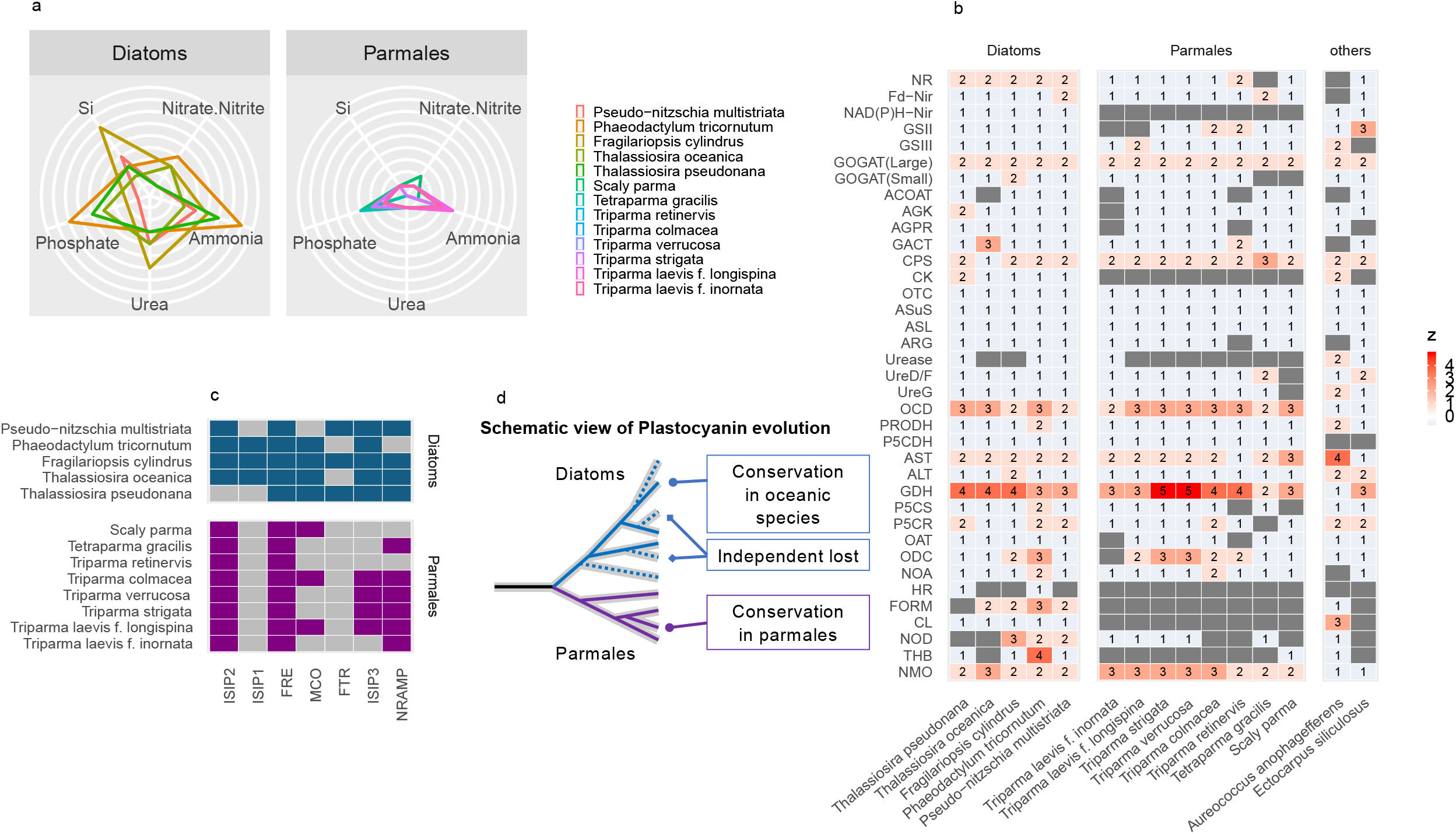
Ecophysiology of diatoms and parmaleans. (a) Distribution of nutrient transporter genes. Each axis represents the number of nitrate/nitrite transporter, ammonia transporter, urea transporter, phosphate transporter, or silicic acid (Si) transporter genes. (b) Genes involved in nitrogen assimilation (including ornithine–urea cycle). The colours are scaled in ascending order from blue to red by the z-value in each row; a grey square indicates absence of the gene. (c) Presence (filled square) or absence/loss (grey square) of iron uptake system genes. Gene names are abbreviated; full names and accessions can be found in Supplementary Data 4. (d) Schematic view of the evolutionary pattern of plastocyanin genes. A whole phylogenetic tree is shown in Supplementary Fig. 3.

Parmaleans had all of the ornithine–urea cycle genes, as with diatoms^15^ and other stramenopiles^41^ (Fig. 3b, Supplementary Fig. 2). Other involved genes (i.e., those encoding NAD(P)H nitrite reductase, carbamate kinase, formamidase, cyanate lyase, and hydroxylamine reductase) were present in diatoms but absent from parmaleans. NAD(P)H nitrite reductase is a major enzyme in nitrogen metabolism that catalyses production of ammonia from nitrite. Carbamate kinase is a major enzyme that produces carbamoyl phosphate, which is a precursor of the urea cycle. In contrast to this difference, both parmaleans and diatoms encode alternative enzymes for these proteins, namely ferredoxin-nitrite reductase and carbamoyl phosphate synthetase, having the same functions as NAD(P)H nitrite reductase and carbamate kinase, respectively. Therefore, parmalean nitrogen metabolism likely relies on the latter set of enzymes, while diatoms may have more efficient nitrogen metabolism by possessing multiple sets of enzymes. Formamidase, cyanate lyase, and hydroxylamine reductase function around the main pathway of nitrogen metabolism. Previous studies showed that formamidase and cyanate lyase are upregulated under N-limited conditions in *Aureococcus anophagefferens*^42^ and *P. tricornutum*^43^. Diatoms encoding these enzymes may have the ability to obtain ammonia from intercellular nitrogen compounds even when they cannot obtain extracellular nitrogen^42,43^. By contrast, parmaleans lacking these enzymes may not have this capacity.

### Iron metabolism

Iron acts as an electron carrier in the photosynthesis system and various metabolic processes in phototrophs. In marine ecosystems, iron is one of the prime limiting elements for phototrophs because of high demand^44^. Therefore, iron uptake ability is an important factor for competition in marine environments. We searched for iron metabolism-related genes in diatom and parmalean genomes. Ferric reductase (FRE), a high-affinity reductive iron uptake system component, was found in all diatoms and parmaleans investigated (Fig. 3c), but parmaleans completely lacked Fe^3+^ permease (FTR) genes (Fig. 3c). Parmalean genomes encoded genes with high sequence similarity to diatom FTR genes, but the parmalean sequences lacked the [REXXE] motif, which is important for iron permeation^45^. This indicates that the diatom/Parmales common ancestor possessed FTR but parmalean FTR homologs may have lost their ability to enable iron permeation during evolution. As for the genes involved in the non-reductive iron uptake system, iron starvation-induced protein 2 (ISIP2)^46^ was widely distributed in parmaleans, whereas ISIP1 was not present (Fig. 3c). ISIP1 plays an important role in siderophore uptake in diatoms and is considered a highly efficient iron uptake gene^47^. Our results support the idea that ISIP1 is a diatom-specific gene^47^ and its presence may underlie diatoms’ high iron uptake capacity.

Most parmaleans encode genes for plastocyanin, a copper-containing redox protein that can substitute for cytochrome *c*_*6*_, which is a redox protein that requires iron and transfers electrons from the cytochrome *b*_*6*_*–f* complex to photosystem I during photosynthesis. It was generally thought that chlorophyll *c*-containing algae lack plastocyanin, but several pelagic diatoms from different genera (including *Thalassiosira oceanica*) encode plastocyanin and are thought to be adapted to iron-deficient pelagic regions^48,49^. Parmaleans may also have an environment-dependent adaptive strategy to differentially use cytochrome *c*_*6*_ and plastocyanin. Phylogenetic analysis revealed that the plastocyanin genes from diatoms and parmaleans were monophyletic (with dictyochophytes and others), except for *Fragilariopsis kerguelensis*, which was grouped with bacteria (Supplementary Fig. 3). This result is inconsistent with the previously proposed horizontal acquisition of plastocyanin genes in pelagic diatoms^48^. The diatom/Parmales common ancestor likely possessed both cytochrome *c*_*6*_ and plastocyanin, and some diatoms (mostly coastal ones) lost their plastocyanin (Fig. 3d).

### Silicate metabolism

Each parmalean genome contained at most one silicic acid transporter (SIT) gene, whereas diatom genomes contained multiple SIT genes (Fig. 3a). Most SIT genes of diatoms and ‘Scaly parma’ encode a 10-fold transmembrane type (i.e., single SIT domain), whereas the SIT genes of the other parmaleans encode a 20-fold transmembrane type (i.e., two SIT domains). Phylogenetic analysis of SIT domains indicated that parmalean SIT genes belong to the most basal clade of diatom SITs (clade B)^50^, and that the 20-fold transmembrane-type SITs of Parmales are the result of multiple (likely two times) domain duplications in the *Triparma* lineage (Fig. 4). A large number of paralogous SITs (at least five clades) in diatoms was generated through multiple gene duplications in the diatom lineage after it diverged from the Parmales lineage.

**Fig. 4.**
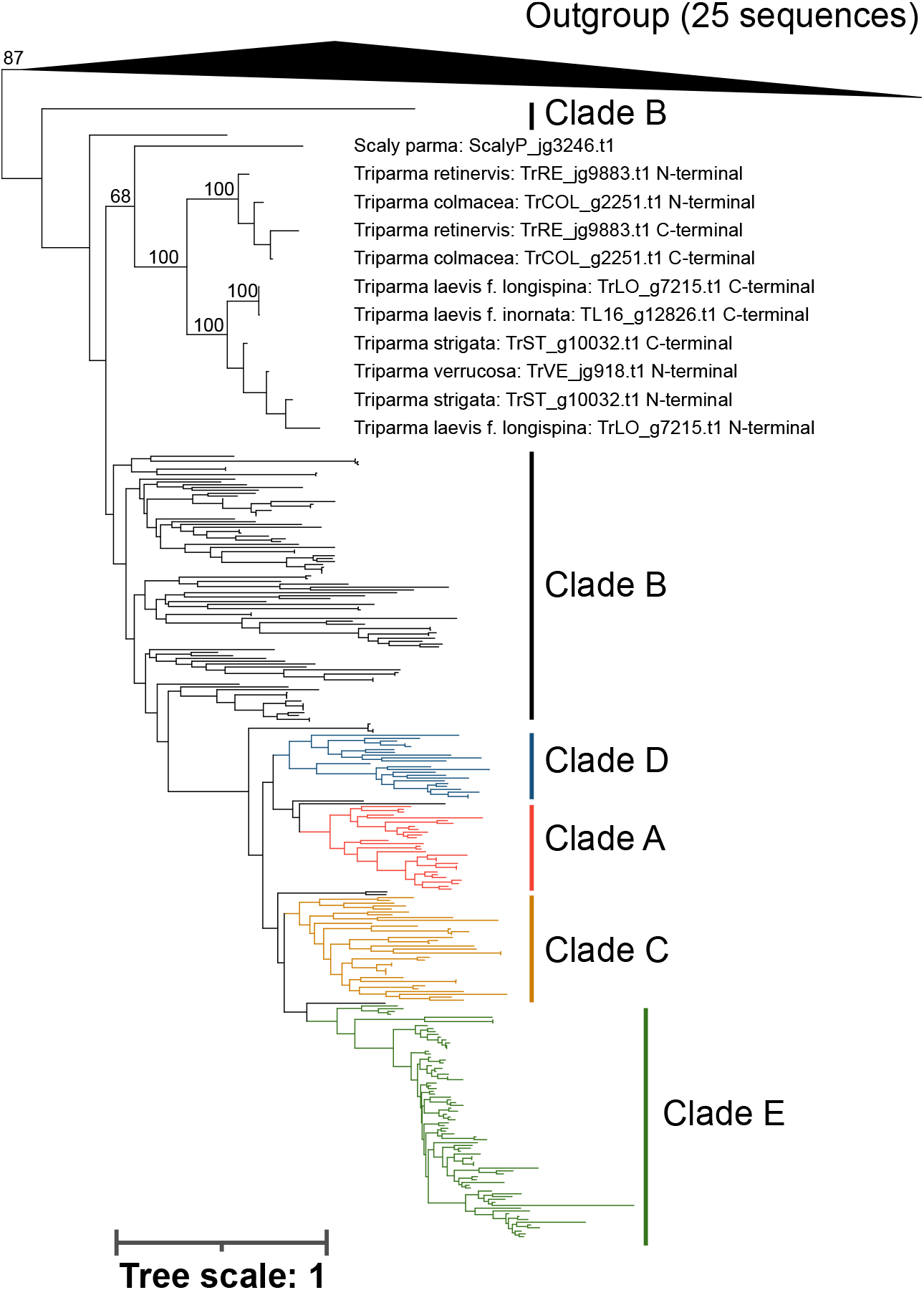
Phylogenetic tree of SIT domains. Maximum likelihood phylogenetic tree of the SIT domains of diatoms, Parmales, and ochrophytes (outgroup). Sequences with more than two SIT domains were separated to each domain and aligned. Grouping of paralogues from diatoms is based on the classification of Durkin et al. (2016). Only important bootstrap values are noted.

Silicanin homologs, some of which are biosilica-associated proteins^51^, were found in Parmales. The parmalean genomes encoded 1–2 silicanin homologs, whereas the diatom genomes encoded 7–14 homologs. Parmalean silcanin homologs have the RXL domain, which is typical of many diatom biosilica-associated proteins^52–55^ but lack the NQ-rich domain that is found in Sin1 and Sin2 of *Thalassosira pseudonana*^51^. Silicanin homologs are not known in other stramenopiles, but have been reported in transcriptome data of other non-diatom eukaryotes such as the ciliate *Tiarina fusus* and the dinoflagellate *Rhizochromulina marina*^51^. These silicanin homologs were found to be highly similar to those of the diatoms *Synedropsis recta* and *Thalassiosira weissflogii*, respectively (blastp search against the MMETSP database; identities are 68.3% and 100%, respectively, and e-values are both 0.0), implying that these genes from non-diatom eukaryotes originated from diatoms (either through HGT or contamination). Thus, the silicanin homologs found in parmaleans are the first examples of non-horizontally transferred silicanin homologs in non-diatom species, implying that the silicanin gene was already present in the diatom/Parmales common ancestor. Silicanins, like SITs, have undergone multiple gene duplications within the diatom lineage after the diatom/Parmales divergence. Interestingly, SIT and silicanin proteins were not found in any bolidomonad transcriptomes, consistent with their lack of silica plates.

### Ecological strategies and evolutionary scenarios

By comparing the genomes of eight parmaleans and five diatoms, we were able to delineate differences and similarities in gene content between these two taxa (Fig. 5). Based on the gene-based trophic model, our analysis suggests that parmaleans are phago-mixotrophs that can acquire nutritional resources such as carbon, nitrogen, phosphorus, vitamins, and trace elements (e.g., iron) by grazing other organisms, such as bacteria. Although phago-mixotrophs would be less dependent on inorganic nutrient resources, this advantage is traded off with an associated increase in metabolic costs for incorporating and maintaining the cellular components required for both autotrophy and phagotrophy. In addition, since phagotrophy reduces the cell surface area for transporter sites, phago-mixotrophs are thought to have lower growth efficiency relative to photoautotrophic specialists^56,57^. According to a theoretical study, mixotrophy is beneficial especially in oligotrophic water, whereas autotrophy is advantageous in eutrophic environments^58,59^.

**Fig. 5.**
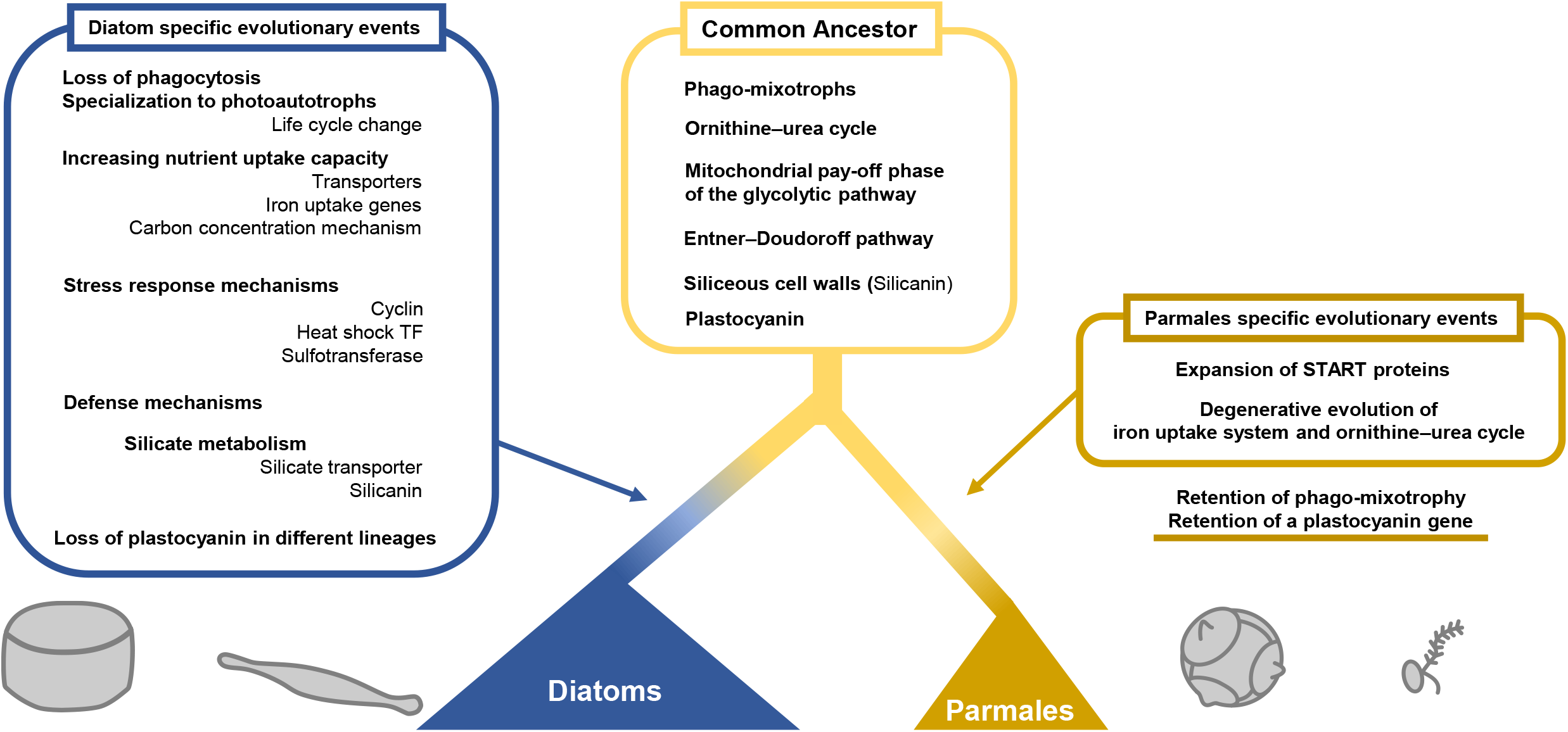
Schematic view of diatoms and parmales evolution.

Previous studies suggested that some mixotrophs can widen their niche by alternating their trophic strategies^60,61^. For example, several coccolithophores (Haptophyta) are known to alternate between a motile phago-mixotrophic haploid stage and a non-motile autotrophic diploid stage based on nutrient condition^62^. Based on these facts and other field data, it has been previously hypothesized that parmaleans have a similar life stage alternation^3^. Namely, parmaleans may live as silicified photoautotrophs during winter (the cold mixing season) when nutrients are rich, while they may feed on bacteria through phagocytosis as naked flagellates during summer (the warm stratified season) when nutrients are depleted. Our study reinforces the possibility of such a life cycle in Parmales, by detecting the genes for phagocytosis which has a potential association with the naked-flagellate stage.

In addition to the absence of phagotrophy in diatoms, our analysis revealed a marked contrast in the gene repertoires between diatoms and parmaleans, with all indicating the autotrophic adaptations of diatoms. For example, there is a large difference in the number of nutrient transporter genes between diatoms and parmaleans (Fig. 3a), clearly representing an adaptation of diatoms to eutrophic environments, although it is not clear whether these paralogous genes have different functions (e.g., affinity, transport rates, and subcellular localization) or a dosage effect^63^. In addition, there are differences in the number of genes involved in biophysical CCMs (Supplementary Fig. 4; See Supplementary Note). Diatoms possess higher CO_2_ fixation capacity relative to other phytoplankton groups^64^, and these gene repertoires may support this trait. We also revealed the expansion of protease and sulfotransferase genes in diatoms in addition to the previously described expansion of cyclin and heat-shock protein genes^16,17^ (Fig. 2a). These genes are likely involved in stress response and population control, which support the extraordinary growth capacity of diatoms.

Most diatoms are photoautotrophs, and all diatoms that we studied were predicted as such (Fig. 2d, e). However, phagotrophy must have existed for the ancestor of diatoms to take up red algae as endosymbionts (i.e., secondary plastids). Some members of ochrophyte, such as chrysophytes and dictyochophytes^65^, are known to be phago-mixotrophic. Our results suggest that Parmales, which is the closest group to diatoms, is also phago-mixotrophic. Thus, the diatom/Parmales common ancestor was firmly inferred as phago-mixotrophic, and there were massive serial gene loss events related to phagocytosis loss and specialization to photoautotrophy in the early evolution of diatoms after diverging from Parmales (ca. 180–240 million years ago^66^).

Diatoms always have silicified cell walls in the vegetative stage, whereas parmaleans putatively switch between two life stages, silicified/non-flagellated and naked/flagellated stages. The silicified cell wall provides a barrier against grazers, parasites, and pathogens^67^, but is obviously incompatible with phagocytosis as it completely covers the cell. Thus, there is a trade-off between silicification/autotrophy and phagocytosis, and the loss of phagotrophy in diatoms may have been related to benefits from the silicified cell wall. To reveal why photoautotrophic diatoms diverged from the phago-mixotrophic lineage and specialized to the silicified life stage, it is necessary to understand not only the costs and benefits associated with mixotrophy but also those of defence by silicified cell walls.

The next possible step in the evolution of diatoms after specialization to silicification and photoautotrophy might have been to thicken their silicified cell wall and increase their cell size^8^. Diatoms tend to have larger cell sizes than parmaleans, and the evolution of these traits has the great advantage of increasing resistance to grazers. The evolution of silicic acid transporter genes (Fig. 4) may have supported the evolution of silicified cell walls because diatoms with thick walls and large cells require large amounts of silicate. It is also known that nutrient metabolism, especially nitrogen metabolism, is closely related to silica deposition in diatoms^68^. Thus, the ability of diatoms to take up nutrients may also be related to the evolution of their silicified cell wall. Silicanin, which diversified in diatoms, may have been important in the precise control of the formation of thick cell walls. It has been also pointed out that vacuoles play a major role in cell size expansion^8^. However, there is little evidence of differences in vacuole-related genes between parmaleans and diatoms (e.g., lack of a vacuolar nitrate transporter homolog in parmaleans), so further discovery and analysis of the relevant genes are needed to address this issue.

Diatoms are also an important group in iron usage in the ocean, often dominating iron-stimulated blooms^69^. Analyses of iron utilization strategies revealed that the ISIP1 gene, which is involved in siderophore-mediated iron acquisition, is absent in parmaleans and unique to diatoms (Fig. 3c). Siderophores are thought to be major components of microbial iron cycling in the ocean^70^. The lack of the ISIP1 gene in parmaleans supports the idea that this gene underlies the high iron uptake capacity of diatoms and supports their photoautotrophic lifestyle. We also found that plastocyanin, which is an alternative for iron-requiring proteins in photosynthesis, is widely distributed in parmaleans. Phylogenetic analysis suggests that each lineage of diatoms lost their plastocyanin genes independently, and that pelagic diatoms and parmaleans conserved plastocyanin genes from their common ancestor (Fig. 3d). Parmaleans retained plastocyanin to balance their restricted capacity for iron uptake in iron-limited environments; diatoms increased their iron uptake capacity (e.g., ISIP1), while several lineages have specialized to coastal eutrophic environments and lost plastocyanin.

Our analysis also revealed that the ornithine–urea cycle, the mitochondrial pay-off phase of the glycolytic pathway, and the Entner–Doudoroff pathway, which have been cited as unique features of diatoms, were substantially conserved from common ancestor of Parmales and diatoms (Fig. 3b, Fig. 5, Supplementary Fig. 2, Supplementary Fig. 5: see Supplementary Note). We also found the expansion of genes related to lipid metabolism and intracellular signalling, and the degenerative evolution of several genes related to iron uptake and ornithine–urea metabolism in Parmales (see Supplementary Note). However, their physiological functions and evolutionary significances remain unclear. Future studies based on a larger set of genomic data will further enhance understanding of the physiology, ecology, and evolution of these fascinating organisms.

## Methods

### Culture

We used strains of the parmaleans *Triparma laevis* f. *inornata* (NEIS-2656), *Triparma laevis* f. *longispina* (NIES-3700), *Triparma verrucosa* (NIES-3699), and *Triparma strigata* (NIES-3701), isolated from the Oyashio region of the western North Pacific. For the other strains, water samples were collected at 10 m in the Notoro-ko lagoon (44º3’2.1” N, 144º9’38.8” E, December 2015) for *Triparma retinervis*, at 10 m in the Sea of Okhotsk (45º25’0” N, 145º10’0” E, June 2017) for *Tetraparma gracilis* and *Triparma columacea*, and at 30 m in the Sea of Okhotsk (44º30’0” N, 144º20’0” E, June 2014) for the uncharacterized ‘Scaly parma’. The strains were isolated by serial dilution with siliceous cell wall labelling techniques described previously^5^. The strains were cultured in f/2 medium^71^ at 5 °C under a light intensity of ca. 30 μmol photons m^− 2^ s^− 1^ (14:10 L:D cycle).

### Genomic DNA, RNA extraction and sequencing

Cells grown under exponential growth phase were harvested by centrifugation, and either DNA (all strains, except *Triparma laevis* f. *inornata*) or RNA (for *Triparma laevis* f. *inornate, Triparma strigata, Triparma retinervis* and ‘Scaly parma’) was extracted using the DNeasy Plant Mini Kit or RNeasy Plant Mini Kit (Qiagen, Venlo, Netherlands), respectively. Libraries were generated using the Illumina TruSeq DNA/RNA sample preparation kit (Illumina, Inc., San Diego, USA). Sequencing of whole genomes or transcriptomes was performed on an Illumina HiSeq X (150 bp, paired-end) or HiSeq 2000 (100 bp, paired-end), respectively. Exceptionally, the genome of *Triparma laevis* f. *longispina* and ‘Scaly parma’ was sequenced with an Illumina HiSeq 2500 (150 bp, paired-end). DNA extraction and sequencing methods for *Triparma laevis* f. *inornate* were already reported in Kuwata et al^13^.

### Genome assembly and microbial sequence contamination removal

Genome assembly and contamination removal methods for *Triparma laevis* f. *inornate* were already reported in Kuwata et al^13^. For the other strains, the Illumina reads were trimmed with Trimmomatic (v.0.38)^72^ using the following parameters: LEADING:20 TRAILING:20 SLIDINGWINDOW:4:15 MINLEN:36 TOPHRED33. The filtered reads were assembled by Platanus (v.1.2.4)^73^ with default options. To remove bacterial contamination from contigs, clustering of contigs was performed based on the coverage calculated by read mapping, GC frequency, and k-mer frequency. In addition, the phylogenetic classification of the genes in the contigs was estimated using lowest common ancestor analysis. The results were used to determine the clusters composed of bacterial contigs. Read mapping to assembled contigs with the filtered reads was performed with BWA (v.0.7.17)^74^. The coverage was calculated from the resulting .sam file using a custom-written Perl script, which was also used to determine the GC content. The tetramer frequency of contigs was calculated using cgat (v.0.2.6)^75^. Open reading frames (ORFs) were predicted using GeneMarkS (v.4.30)^76^ and their taxonomy was annotated with a last common ancestor strategy as in Carradec et al^77^. ORFs were searched against a database composed of UniRef 90^78^, MMETSP database^79^, and Virus-Host DB^80^ using DIAMOND (v.0.9.18)^81^. Selected hits were then used to derive the last common ancestor of the query ORFs with the NCBI taxonomy database. Clustering of contigs was performed using the R script provided in the CoMet workflow^82^ with coverage, GC content, and k-mer frequency as information sources. The organism from which each cluster originated was determined from the estimated phylogeny of the genes in the contigs belonging to the cluster. Contigs belonging to bacterial-derived clusters were excluded from the datasets and not used in downstream analyses.

We also performed a blastn (v.2.11.0) search against the organelle genomes of *Triparma laevis* f. *inornata*^83^ to remove the organelle genome from assembled contigs. Contigs that hit the organelle genome of *T. laevis* f. *inornata* with E-values < 1e− 40 were excluded from our dataset as organelle genomes.

### Genome annotations

For *T. laevis* f. *inornata* genome^13^, rRNA and tRNA genes were predicted by Barrnap (v.0.6, http://www.vicbioinformatics.com/software.barrnap.shtml) and tRNA-scan-SE (v.1.23)^84^, respectively. The protein coding-genes were predicted by AUGUSTUS (v.3.2.2)^85^ with the RNA-seq data mentioned above. First, the RNA-seq reads processed by fastx-toolkit (v.0.0.13, http://hannonlab.cshl.edu/fastx_toolkit/) were mapped to the contig of *T. laevis* f. *inornata* nuclear genome and assembled into transcript contigs using Tophat (v.2.1.1)^86^, Cufflinks (v.2.2.1)^87^ and Trinity (v.2.0.6)^88^, respectively. The diatom protein sequences from *Thalassiosira pseudonana*^15^ and *Phaeodactylum tricornutum*^16^ were subsequently aligned to the transcript contigs using tblastn search (v.2.2.29) and Exonerate (v.2.4.0)^89^ for detecting CDS regions in the *T. laevis* f. *inornata* genome. Finally, a total of 687 loci on the *T. laevis* f. *inornata* contigs were selected as those carrying full-length CDSs and used for parameter fitting in training hidden Markov models in AUGUSTUS. In gene prediction, the mapping data from both RNA-seq reads and diatom protein sequences were utilized as hints in AUGUSTUS.

For other genomes, tRNA genes were predicted using tRNAscan-SE (v.2.0.7)^90^. Non-coding RNAs excluding tRNAs but including rRNAs were predicted with the Rfam database using infernal (v.1.1.3)^91^. Repeats and transposable elements were annotated and soft-masked using RepeatModeler (v.2.0.1)^92^ and RepeatMasker (v.4.1.0)^93^. For *Triparma strigata, Triparma retinervis* and ‘Scaly parma’, the protein-coding genes were predicted by BRAKER2^94^ with the RNA-seq data mentioned above and a reference protein sequence database. We generated a reference protein sequence database for BRAKER2 from OrthoDB^95^, MMETSP database^79^ and *T. laevis* f. *inornata* protein sequences predicted previously. Firstly, RNA-seq data were mapped to the contigs by STAR (v.2.7.3a)^96^, generating a .bam file. Secondly, BRAKER2 was run in – etpmode mode with the generated .bam file and the reference protein sequence database as the protein hints. For *Triparma laevis* f. *longispina, Triparma verrucosa, Triparma columacea* and *Tetraparma gracilis*, the protein-coding genes were predicted by BRAKER2 only with a reference protein sequence database. We updated the mentioned reference protein sequence database with the predicted protein sequences form *Triparma strigata, Triparma retinervis*, and ‘Scaly parma’, and generated a new database. Finally, BRAKER2 was run in -epmode using the newly generated reference protein sequence database as the protein hints.

### Functional Annotation

For methodological consistency, we applied the same annotation pipelines for our novel genomes and the genomes downloaded from public databases. Genes were functionally annotated by InterProScan (v.5.26-65.0)^97^ and eggNOG-Mapper (v.2.0.1)^98^ with the eggNOG database (v.5.0)^99^. Protein localization was predicted using MitoFates (v.1.1)^100^, TargetP (v.2.0)^101^, SignalP (v.4.1)^102^, and ASAFIND (v.1.1.7)^103^. Protein functions and localizations were manually curated for detailed analyses.

### Phylogenomic analysis

Orthologous genes (OGs) were determined by OrthoFinder (v.2.3.7)^104^ with protein sequences of parmalean genomes, other available stramenopile genomes, and ochrophyte transcriptomes (Supplementary Data 11) from the MMETSP database^79^. Only single-copy genes in each OG and genes that were found in the 18 stramenopile species were retained for downstream phylogenomic analysis, resulting in 164 OGs. Gene sequences within each OG were aligned using MAFFT (v.7.453)^105^ in the linsi mode, and poorly aligned regions from the multiple sequence alignment were removed by trimAl (v.1.4.1)^106^ in the automated1 mode. The resulting supermatrix contained 50,707 amino acid positions for 18 species, with 5.85% missing data. A maximum likelihood tree was inferred by RAxML (v.8.2.12)^107^ with the partition information of each gene and the LG + F model. We performed 1,000 bootstrap replicates and all bootstrap values were 100, indicating full support.

### Gene family and protein domain analysis

Significant differences in protein domain content annotated by InterProScan between the compared genomes were identified using Fisher’s exact test to calculate the *p-value* for the difference in the number of InterPro domains between parmalean and diatom genomes. The *p-values* were corrected for multiple comparisons using Bonferroni correction. For each genome, protein sequences with 100% similarity to other genes were removed using CD-HIT (v.4.8.1)^108^ with the parameters -c 1 -aS 1.

### Predictions of phago-mixotrophy using a gene-based model

Predicted protein data from eight parmalean genomes and five diatom genomes were tested for phagocytotic potential using a gene-based model described by Burns et al^27^. To determine the phagocytotic potential of parmaleans, we also tested the five transcriptomes of the naked flagellate (bolidomonads) from the MMETSP database^79^ and Kessenich et al. (2014)^109^. Gene annotation was not available for the data from Kessenich et al. (2014)^109^; therefore, coding sequences were annotated using TransDecoder (v.5.5.0) (https://github.com/TransDecoder/TransDecoder).

### Phylogenetic analysis of silicon transporter domains

We used the sequence data of SIT proteins of diatoms and ochrophytes provided by Durkin et al. (2016)^50^ in addition to those of parmaleans determined in this study. Diatom and parmalean SIT proteins are usually composed of a single SIT domain, but some contain more than two domains. To analyse multiple domains at once, SIT domain regions were determined using HMMER (v.3.3.2) with PF03842 from using the profile’s GA gathering cutoff (--cut_ga mode) and selected for downstream analysis. Each SIT domain sequence was aligned using MAFFT (v.7.453) in the linsi mode and unreliable sequences were manually removed. A maximum likelihood phylogenetic tree was inferred from this multiple alignment using RAxML (v.8.2.12). The amino acid substitution model was automatically determined to be the LG model by the software. Bootstrap values were obtained based on 100 bootstrap replicates.

### Phylogenetic analysis of plastocyanin

We used HmmerSearch (v.3.3) with TIGR02656.1 from TIGERFAMs using the profile’s GA gathering cutoff (--cut_ga mode) to find plastocyanin genes in Uniref 90^78^, MMETSP^79^, and our genomes. The genes from MMETSP clustered with 97% similarity using CD-HIT (v.4.8.1). Unreliable sequences were removed manually. We next obtained 716 plastocyanin genes of photosynthetic eukaryotes, cyanobacteria, and cyanophages. Because of the large divergence of the sequences and small number of alignable regions, we used gs2, a software to conduct the Graph Splitting (GS) method^110^, which can resolve the early evolution of protein families using a graph-based approach, to estimate the phylogenetic tree of plastocyanin. We ran the GS method with 100 replicates using the Edge Perturbation method for statistically evaluating branch reliability.

## Data availability

Sequence data generated during the current study are available in DDBJ bioprojects, under accession number PRJDB14101 (RNA reads for *Triparma laevis* f. *inornata*), PRJDB13844 (DNA reads for the other seven strains), and PRJDB13933 (RNA reads for the other seven strains). The assembly data analysed during the current study are also available in the DDBJ repository, under accession numbers BLQM01000001-BLQM01000902 (*Triparma laevis* f. *inornata*), BRXW01000001-BRXW01001055 (*Triparma laevis* f. *longispina*), BRXX01000001-BRXX01000659 (*Triparma verrucosa*), BRXY01000001-BRXY01000634 (*Triparma strigata*), BRXZ01000001-BRXZ01008760 (*Triparma retinervis*), BRYA01000001-BRYA01001858 (*Triparma colmacea*), BRYB01000001-BRYB01007082 (*Tetraparma gracilis*), and BRYC01000001-BRYC01001921 (‘Scaly parma’). Data underlying Figs. and Supplementary Figs. are provided as Supplementary Data files.

## Author Contributions

H.B. performed most of the bioinformatics analyses presented in this work and wrote initial version of the manuscript. R.B.-M., H.E., and H.O. supervised the bioinformatics part of the study. Y.N. performed genome assembly and gene prediction for *Triparma laevis* f. *inornata*. A.K. coordinated the genome sequencing part of the study. S.S., S.Y., K.Y., M.I., and A.K. contributed to culture and DNA/RNA sequencing. N.S. contributed to functional interpretation of the genomes. All authors contributed to the interpretation of the results and the finalization of the manuscript.

## Competing interests statement

The authors declare no competing interests.

## Acknowledgements

This work was supported by JSPS/KAKENHI (No. 22657027, 23370046, 26291085, 221S0002, 16K07489, 16H06279, 17H03724), the Canon Foundation, the Collaborative Research Program of Institute for Chemical Research, Kyoto University (No. 2016-30, 2015-39), and the JST “Establishment of University Fellowships Towards The Creation of Science Technology Innovation” Grant Number JPMJFS2123. Computational time was provided by the SuperComputer System, Institute for Chemical Research, Kyoto University. We thank Gabe Yedid, Ph.D., from Edanz (https://jp.edanz.com/english-editing-b) for editing a draft of this manuscript.

## Notes

### Competing Interest Statement

The authors have declared no competing interest.

