## Supplementary information for "Genome analysis of Parmales, a sister group of diatoms, reveals the evolutionary specialization of diatoms from phago-mixotrophs to photoautotrophs"

**This PDF file includes:**

Supplementary Note

Supplementary Fig. 1 to 5

Captions for Supplementary Data 1 to 11

### Supplementary Note

**No clear evidence of horizontal gene transfers in nitorogen metabolism genes**

Previous studies have suggested that some of the genes involved in diatom nitrogen metabolism, such as carbamate kinase and NADP(H) nitrite reductase, originated from bacteria through horizontal gene transfer (HGT)[1–3]. Such HGTs in diatoms may account for the absence of these genes in parmaleans. However, previous molecular phylogenetic analyses[4,5] indicate another possibility. The NAD(P)H nitrite reductase tree shows monophyly of diatoms and other ochrophytes, indicating that NAD(P)H nitrite reductase was vertically inherited by diatoms from ancestral ochrophytes but lost in parmaleans. In the cases of carbamate kinase, formamidase, cyanate lyase and hydroxylamine reductase, diatom sequences cluster with other eukaryotes, particularly dinoflagellates and haptophytes that contain secondary plastids derived from red algal endosymbionts, and Archaeplastida. This indicates the possibility that diatoms acquired these genes from their secondary symbiont and parmaleans lost them.

**Carbon metabolism**

Aquatic phototrophs, including diatoms and parmaleans, require $CO_2$ for photosynthesis. However, the concentration of $CO_2$ dissolved in water is much lower than the required concentration; therefore, they take up dissolved inorganic carbon (both $CO_2$ and $HCO_3^-$) from water and concentrate it in their cells to increase the $CO_2$ concentration around Rubisco and efficiently fix carbon[6]. These carbon concentrating mechanisms (CCMs) are called biophysical CCMs and are distinguished from biochemical CCMs, as found in terrestrial C4 plants. There are two main types of proteins involved in biophysical CCMs[7–9]: carbonic anhydrases (CA), of which there are several classes[10], and bicarbonate ($HCO_3^-$) transporters. In our results, each diatom genome contained about 20 CAs, whereas the parmalean genomes contained fewer than 10 CAs (Supplementary Fig. 4a). Of the seven classes of CAs present in the diatom genomes, the α- and ι-classes were absent from the parmalean genomes. By contrast, β-class CAs were present in all parmaleans but absent in all diatoms except *Phaeodactylum tricornutum*. Regarding bicarbonate transporters, 7–15 genes were present in the diatom genomes, whereas 5–7 genes were present in the parmalean genomes (Supplementary Fig. 4b). Thus, diatoms have more genes involved in CCA than do parmaleans.

Glycolysis, gluconeogenesis, and pyruvate hub metabolism[11] are central pathways of carbon metabolism in diatoms. Previous studies have shown that diatoms have more genes involved in carbon metabolism than do green algae[11,12]. However, we found no significant differences in the number of genes involved in carbon metabolism and the predicted localization of gene products between diatoms and parmaleans (Supplementary Fig. 5). The mitochondrial pay-off phase of glycolysis, which is a known feature of diatom carbon metabolism, was also predicted to be present in parmaleans. Furthermore, this localized phase was also found in oomycetes and other ochrophytes in our datasets. Therefore, the mitochondrial pay-off phase is not specific to diatoms but common to stramenopiles.

Fusion genes for TPI–GAPDH[11], which are the first enzymes of the diatom mitochondrial pay-off phase of glycolysis, were present in seven parmalean genomes (missing in *Triparma laevis* f. *inornata*). Excluding *Tetraparma gracilis*, the fusion genes found in parmaleans target the mitochondria, as in diatoms. An additional GAPDH gene located upstream (and on the opposite strand) of the fusion gene conserved in most diatoms was also found in five parmalean genomes. This GAPDH gene was not found in the 'Scaly parma' and *Tetraparma gracilis* genomes, in which the TPI-GAPDH fusion gene was located at the end of the contig. It was previously suggested that this conserved gene order contributes to the coordinated regulation of the two genes using a bidirectional promoter and plays an important role at the starting point of glycolysis[11]. This characteristic gene order probably emerged in the common ancestor of diatoms and Parmales, as it was not found in other ochrophytes (*Aureococcus anophagefferens* and *Ectoculpus siliculosus*).

Diatoms use a prokaryote-like Entner–Doudoroff (ED) pathway for mitochondrial glycolysis[13]. In this pathway, 6-phosphogluconate dehydratase (EDD) and 2-keto-3-deoxyphosphogluconate aldolase (EDA) play major roles. Many stramenopiles possess EDD genes. By contrast, EDA genes are absent in stramenopiles except diatoms. Thus, a previous study suggested that the diatom EDA originated by HGT from bacteria[13]. In the present work, parmaleans were found to possess both EDD and EDA genes (Supplementary Fig. 5). This result indicates that the prokaryote-like ED pathway is not specific to diatoms and was probably acquired from the diatom/Parmales common ancestor.

#### Supplementary Figures

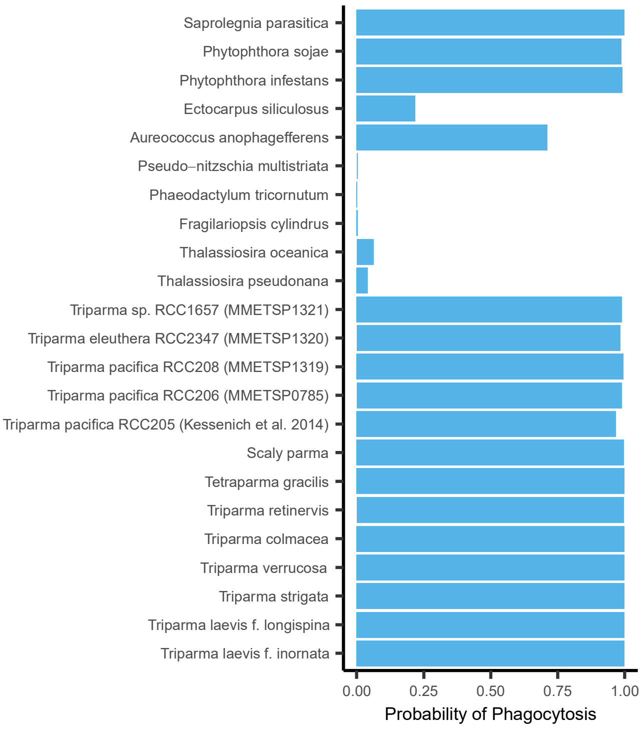


Supplementary Fig. 1 | Probability of phagotrophy predicted by the Burns et al. (2019) tool.

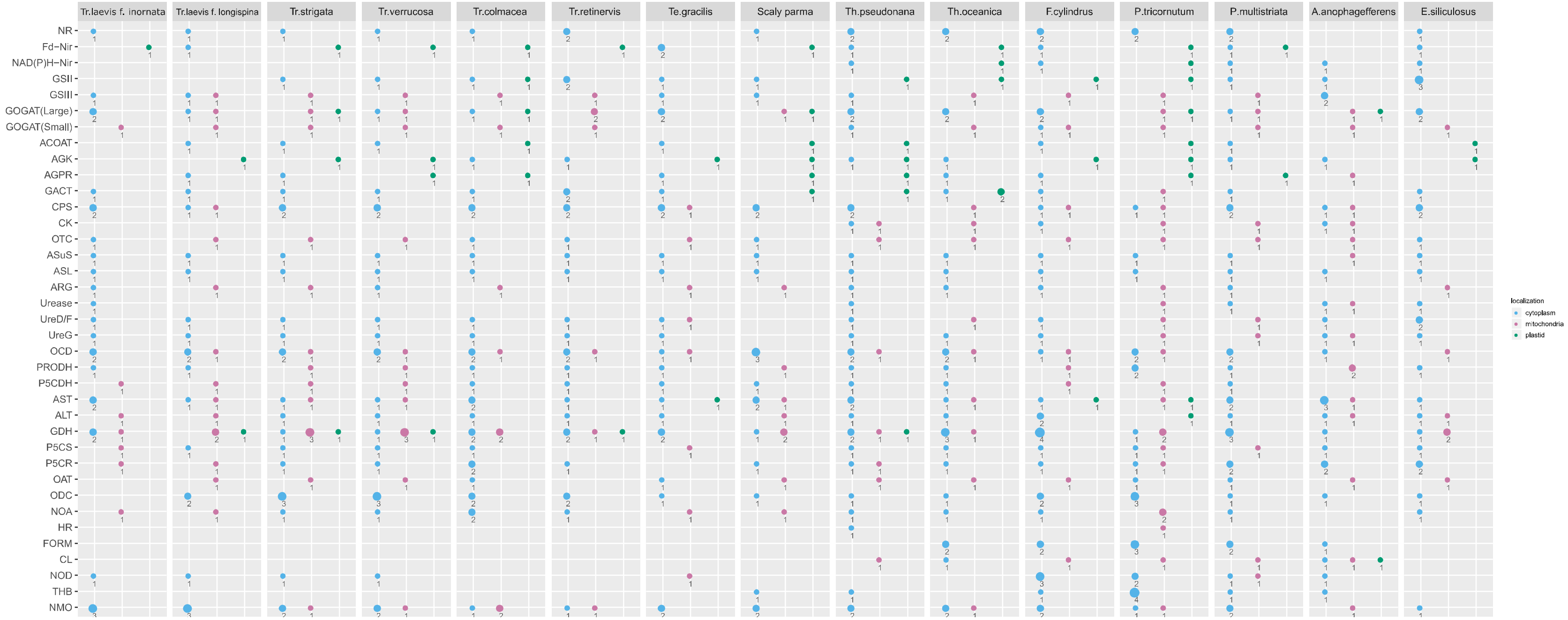


Supplementary Fig. 2 | Genes in nitrogen metabolism and their predicted intercellular localization.

Circle colours represent predicted intercellular localization. Pink: mitochondria; green: chloroplast; blue: cytoplasm. The size of the circle and the number below represent the number of associated genes. Gene names are abbreviated; full names and accessions can be found in Supplementary Data 6.

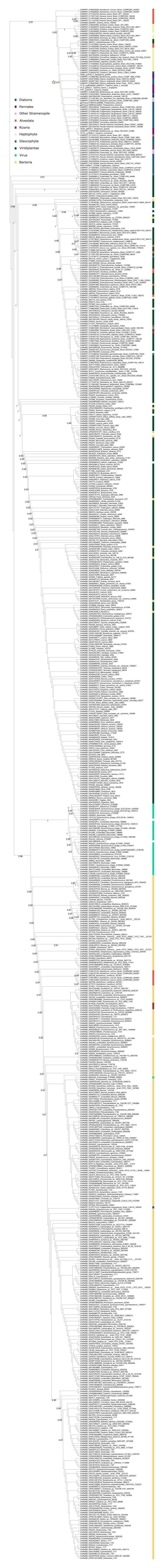
Diatoms
Parmales
Other Stramenopile
Alveolata
Rizaria
Haptophyte
Glaucophyte
Viridiplantae
Virus
Bacteria

Supplementary Fig. 3 | Phylogenetic tree of plastocyanin genes of photosynthetic eukaryotes, cyanobacteria, and cyanophages.

The Graph Splitting method[110] was used to reconstruct the tree, with 100 replicates for the Edge Perturbation (EP) method for statistically evaluating branch reliability. Only EP scores > 0.9 are shown in the figure. The star represents the node supporting the monophyly of plastocyanin genes of diatoms and parmaleans (EP scores = 0.91).

a

| taxa | α | β | γ | δ | ζ | θ | ι | Total |
|---|---|---|---|---|---|---|---|---|
| Pseudo–nitzschia multistriata | 5 | 0 | 4 | 1 | 0 | 2 | 3 | 15 |
| Phaeodactylum tricornutum | 5 | 2 | 4 | 0 | 0 | 4 | 1 | 16 |
| Fragilariopsis cylindrus | 7 | 0 | 3 | 2 | 0 | 6 | 1 | 19 |
| Thalassiosira oceanica | 2 | 0 | 4 | 4 | 1 | 3 | 3 | 17 |
| Thalassiosira pseudonana | 3 | 0 | 4 | 4 | 1 | 5 | 1 | 18 |
| Scaly parma | 0 | 3 | 2 | 0 | 0 | 2 | 0 | 7 |
| Tetraparma gracilis | 0 | 3 | 2 | 0 | 0 | 1 | 0 | 6 |
| Triparma retinervis | 0 | 1 | 2 | 0 | 1 | 2 | 0 | 6 |
| Triparma colmacea | 0 | 1 | 2 | 0 | 0 | 1 | 0 | 4 |
| Triparma verucossa | 0 | 1 | 3 | 1 | 0 | 3 | 0 | 8 |
| Triparma strigata | 0 | 1 | 3 | 1 | 0 | 3 | 0 | 8 |
| Triparma leavis f. longispina | 0 | 1 | 3 | 2 | 0 | 3 | 0 | 9 |
| Triparma leavis f. inornata | 0 | 1 | 3 | 0 | 0 | 1 | 0 | 5 |
| Ectocarpus siliculosus | 1 | 2 | 3 | 0 | 0 | 0 | 0 | 6 |
| Aureococcus anophagefferens | 1 | 0 | 3 | 4 | 0 | 0 | 0 | 8 |

b

| taxa | SLC4 | SLC26 |
|---|---|---|
| Pseudo–nitzschia multistriata | 4 | 5 |
| Phaeodactylum tricornutum | 8 | 4 |
| Fragilariopsis cylindrus | 6 | 6 |
| Thalassiosira oceanica | 6 | 9 |
| Thalassiosira pseudonana | 3 | 4 |
| Scaly parma | 1 | 3 |
| Tetraparma gracilis | 3 | 3 |
| Triparma retinervis | 2 | 5 |
| Triparma colmacea | 2 | 3 |
| Triparma verrucosa | 2 | 4 |
| Triparma strigata | 2 | 5 |
| Triparma laevis f. longispina | 2 | 4 |
| Triparma laevis f. inornata | 2 | 4 |
| Ectocarpus siliculosus | 2 | 2 |
| Aureococcus anophagefferens | 3 | 4 |

Supplementary Fig. 4 | Genes potentially involved in biophysical carbon concentration mechanisms (CCMs).

(a) Number of carbonic anhydrase genes potentially involved in biophysical carbon concentration mechanisms (CCMs). Greek letters (α, β, γ, δ, ζ, θ, ι) indicate the gene families (classes)[111]. (b) Number of bicarbonate transporter genes potentially involved in biophysical CCMs.

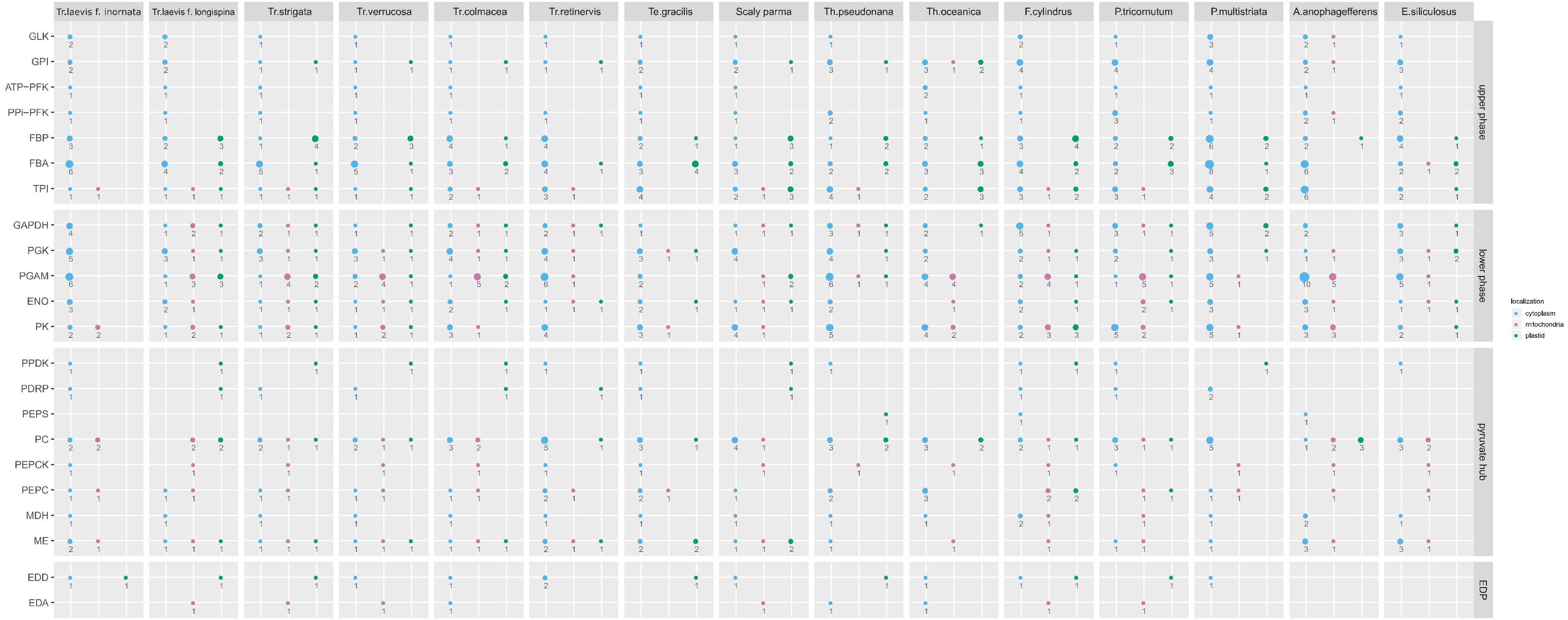


Supplementary Fig. 5 | Genes in carbon metabolism and their predicted intercellular localization.

Circle colours represent predicted intercellular localization. Pink: mitochondria; green: chloroplast; blue: cytoplasm. The size of the circle and the number below represent the number of associated genes. Gene names are abbreviated; full names and accessions can be found in Supplementary Data 10.

#### Description of supplementary files

**Supplementary Data 1**

InterPro domains enriched in diatom genomes. InterPro domain IDs (column A), InterPro domain description (B), Number of genes(C-O), a total number of genes annotated with this domain(P,Q), a total number of genes not annotated with this domain (R,S), *p-values* for Fisher's exact test (T), corrected *p-values* for multiple comparisons by Bonferroni correction (U) and manually assigned categories for InterPro domains(V).

**Supplementary Data 2**

InterPro domains enriched in parmalean genomes. Descriptions for column are same with Supplementary Data 1.

**Supplementary Data 3**

InterPro domains for START protein genes. InterPro domain IDs (column A), InterPro domain description (B), Number of genes(C-Q), a total number of genes annotated with this domain in parmalean genomes(R).

**Supplementary Data 4**

Intraflagellar transport (IFT) subunits gene/transcripts catalog. Gene accessions (column A), gene name(B), InterPro ID / EggNOG ID used to estimate gene function (C), gene category (D) taxonomic name (E), and group names they belong to (F).

**Supplementary Data 5**

Transporter gene catalog. Gene accessions (column A), gene name abbreviation (B), gene name(C), InterPro ID / EggNOG ID used to estimate gene function (D), taxonomic name (E), and group names they belong to (F).

**Supplementary Data 6**

Nitrogen metabolism gene catalog. Gene accessions (column A), gene name abbreviation (B), gene name(C), InterPro ID / EggNOG ID used to estimate gene function (D), taxonomic name (E), group names they belong to (F), assigned subcellular localization based on target predictions (G-L), finally assigned subcellular localization(M), taxonomic name (N) and group names they belong to (O).

**Supplementary Data 7**

Iron metabolism gene catalog. Gene accessions (column A), gene name abbreviation (B), gene name(C), InterPro ID / EggNOG ID used to estimate gene function (D), taxonomic name (E), and group names they belong to (F).

**Supplementary Data 8**

Carbonic anhydrase gene catalog. Gene accessions (column A), gene name(B), InterPro ID / EggNOG ID used to estimate gene function (C), taxonomic name (D), and group names they belong to (E).

**Supplementary Data 9**

Bicarbonate transpoter gene catalog. Gene accessions (column A), gene name(B), InterPro ID / EggNOG ID used to estimate gene function (C), taxonomic name (D), and group names they belong to (E).

**Supplementary Data 10**

Carbon metabolism gene catalog. Gene accessions (column A), gene name abbreviation (B), gene name(C), pathway name (D), InterPro ID / EggNOG ID used to estimate gene function (E), taxonomic name (F), group names they belong to (G), assigned subcellular localization based on target predictions (H-M), finally assigned subcellular localization(N), taxonomic name (O) and group names they belong to (P).

**Supplementary Data 11**

Transcriptome data used in orthologous genes (OGs) clustering.
